# Stress-actuated Flexible Microelectrode Arrays for Activity Recording in 3D Neuronal Cultures

**DOI:** 10.1101/2024.12.12.628189

**Authors:** João Serra, José C. Mateus, Susana Cardoso, João Ventura, Paulo Aguiar, Diana C. Leitao

## Abstract

Microelectrode arrays (MEAs) are instrumental in monitoring electrogenic cell populations, such as neuronal cultures, allowing high precision measurements of electrical activity. Although three-dimensional neuronal cultures replicate the behavior of *in vivo* systems better than two-dimensional models, conventional planar MEAs are not well suited to capture activity within such networks. Novel MEA geometries can overcome this difficulty, but often at the cost of increased fabrication complexity. Here, we used the stress mismatch between thin film layers to fabricate MEAs with vertical electrodes, using methods compatible with established microfabrication protocols. A micrometric SiO_2_ hinge enables control over the bending angle of flexible polyimide structures with embedded electrodes. The performance of the patterned electrodes was assessed before and after stress actuation, through impedance measurements, voltage noise mapping, and neuronal activity recordings. 3D MEAs with 30×30 µm^2^ electrodes showed an impedance of 0.96 ± 0.07 MΩ per electrode and detected neuronal activity spikes with amplitudes as high as 400 µV. These results demonstrate the potential of the developed methods to provide a scalable approach to fabricate 3D MEAs, enabling enhanced recording capabilities for *in vitro* neuronal cultures.

## INTRODUCTION

In electrophysiology, Microelectrode Arrays (MEAs) are the tools of choice for evaluating the generation and transmission of electrical signals in neuronal populations. ^1^ MEAs allow the recording of *in-vitro* neuronal activity with high signal precision^2^ and are capable of electrophysiological stimulation.^3^ They play a fundamental role in the study of neurodegenerative disorders such as Parkinson’s disease^4^ or epilepsy.^5^ Recent developments in MEAs technologies include complementary metal oxide semiconductor (CMOS) integration for high density electrodes,^6^ protruding architectures for enhanced neuron-electrode coupling^7^ and the fabrication of flexible structures for better compatibility with biological tissues.^8^ The use of MEAs to record the activity of a cellular culture was previously limited to the two-dimensional (2D) plane.^9^ However, the 2D models used in most electrophysiology studies fail to adequately replicate the behavior of *in vivo* systems, spurring interest in growing and recording the activity of three-dimensional (3D) neuronal cultures.^10^ This has led to significant developments towards providing tools that are capable of fully capturing the electrical activity of a 3D neuronal network.^11^

The fabrication of devices with 3D geometries has been inspired by origami^12^ and other self-assembling methods. These include strategies such as the use of heat shrinking polymers^13^ or hydrogel swelling.^14^ Another promising approach is to manipulate stress in patterned thin films to achieve specifically tailored shapes, allowing 3D geometries to be extracted from the 2D plane.^15,16^ In cell electrophysiology recording, devices such as stress-actuated cell grippers^17^ or strain-engineered Si probes for cell penetration^18^ have been demonstrated. A significant development was the fabrication of sharp and rigid electrodes that protrude from the substrate due to the combination of a layer with *compressive* stress and a layer with *tensile* stress.^19^ In this case, a SiO_2_/Si_3_N_4_ bilayer was patterned into sharp tips, and the residual stresses led to self-folding electrodes with the ability to penetrate organoids for activity recording. Replacing the traditional rigid materials with flexible layers allows MEAs to more closely match the mechanical properties of biological cells.^20^ Polyimide (PI) is commonly used in microdevices due to its flexibility,^21^ biocompatibility,^22^ and integration with established microfabrication steps.^23^ PI has been used in arrays of electrodes lifted using mechanical actuation.^9^ This solution allows simple fabrication of 3D electrode arrays, but the use of mechanical actuation requires precise manipulation of the individual structures, hindering scaling up.^19^

Here, we harness the stress in thin films to deliver arrays of recording electrodes in vertical structures (Fig. 1). A hinge with specific mechanical properties and dimensions is used to control the bending angle of the structures that contain the electrodes (Fig. 1a). The fabricated devices are shown in Fig. 1b. The protocol used enables reproducible and scalable fabrication of 3D MEAs, allowing seamless access to platforms that can probe the electrical activity of 3D neuronal cultures and study the propagation of neuronal signals in 3D space. The feasibility of our platform is discussed, taking into account the performance of MEAs before and after stress actuation in terms of electrode impedance and voltage noise levels. Validation is performed by recording neuronal activity.

**Figure 1:**
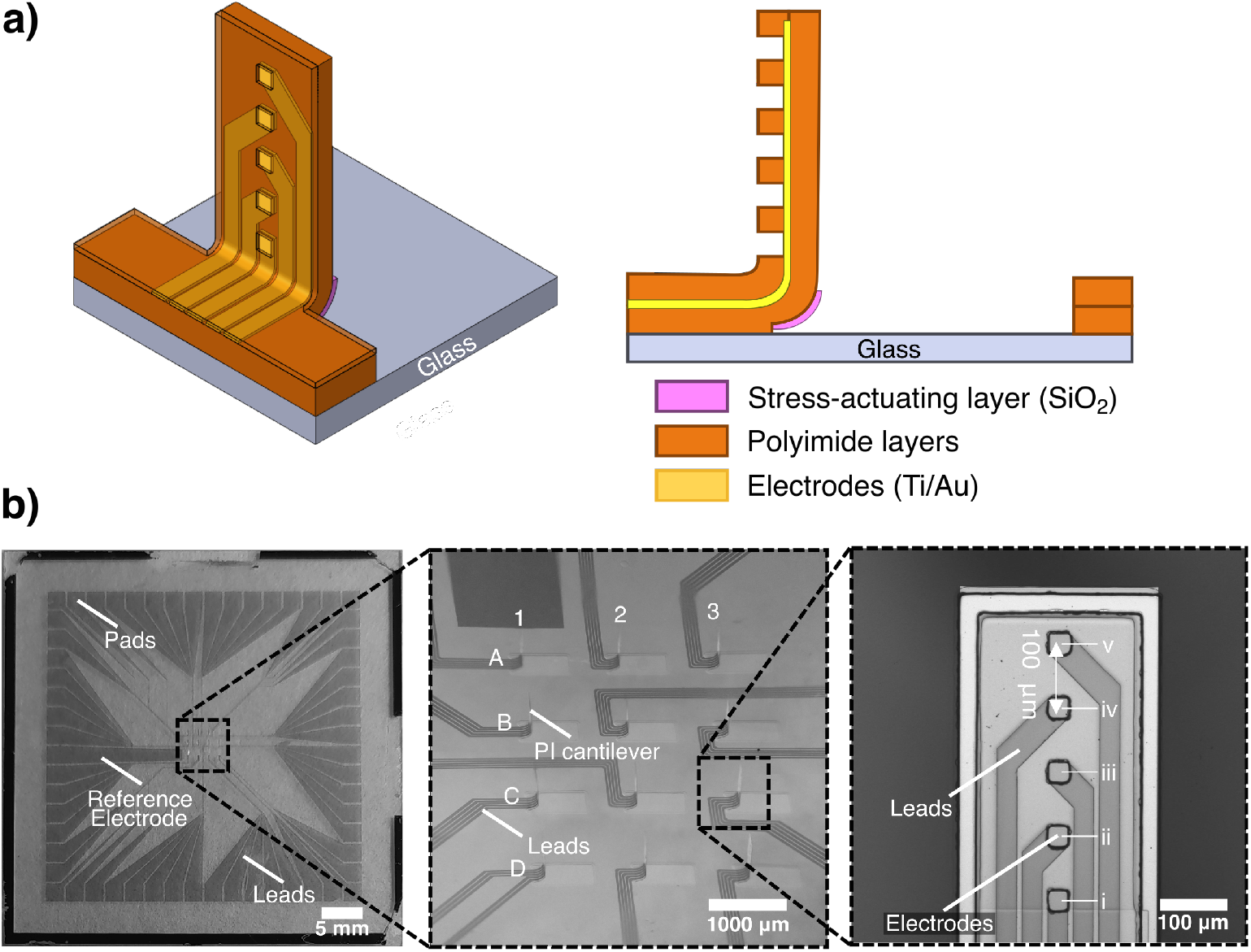
MEAs with 3D recording capabilities based on thin film stress actuation. a) Schematic representation of the designed device, with a stress-inducing SiO_2_ layer forming a hinge which imposes a specific bending angle on the PI cantilever. b) Optical images of a microfabricated 3D MEA: (left) 60-electrode MEA with 3D electrodes obtained with stress actuation, (center) Central electrode area showing the vertical PI cantilevers, and (right) Detail of a PI cantilever containing 5 recording electrodes, taken before bending.

## METHODS

### MEAs fabrication

Planar MEAs were micropatterned following the steps in Fig. 2. The substrate used was glass (Corning Eagle XG) with 49 mm × 49 mm and a thickness of 1.1 mm. First, a sacrificial layer is defined to allow the release of the vertical structures with electrodes. This layer is composed of sputtered Al_98.5_Si_1.0_Cu_0.5_ 150 nm patterned using lithography and wet etching with Technietch Al80 (Microchemicals GmbH). Optical lithography is used in all patterning steps in this work. Afterwards, a SiO_2_ 200 nm thin film is sputter deposited and patterned, which will act as a stress inducing layer to actuate the electrodes. Polyimide (PI-2555 polyimide precursor, HD MicroSystems) was then spin-coated on the substrates at 3 krpm for 30 s, yielding a thickness of about 3.3 µm. The samples were previously covered with a promoter (VM-652, HD MicroSystems) to improve the adhesion of PI to the substrate. PI baking was performed in two steps, first at 120°C for 30 s and then at 140°C also for 30 s. These temperatures were optimized to allow for the transfer of the desired patterns onto the PI using a TMAH-based solution. The PI cantilevers were then defined by lithography, and using TMA238WA for 75 s acted both as a developer of the photoresist and as an etchant of the PI precursor. A final PI curing step was performed by first increasing the temperature to 200 °C at 300 °C/hour, followed by 30 mins at 200 °C and ending with gradual cooling. Ti 5 nm/Au 20 nm electrodes were then defined on top of the PI, grown by sputter deposition and patterned via lift-off. A second PI layer was then used to encapsulate the electrical leads. PI was again spin-coated and patterned as described above, followed by a 30 min curing process at 200 °C. Openings through the PI to the electrodes were then defined to allow electrical contact with the biological medium. Both 3D MEAs and planar MEAs (control samples for performance evaluation) were fabricated and characterized. The 3D MEAs underwent the patterning process shown in Fig. 2 until step G, while the fabrication of planar MEAs was stopped prior to step F.

**Figure 2:**
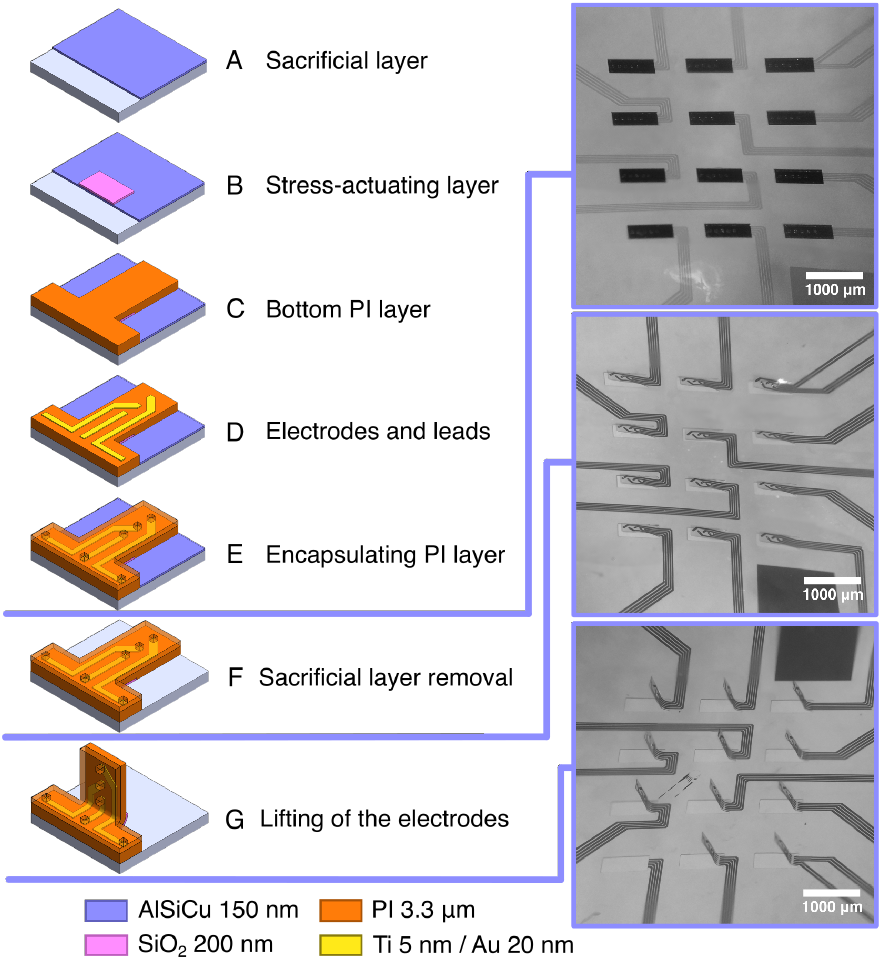
Fabrication process overview for planar-to-3D MEAs based on stress actuation highlighting (left) the major steps of the patterning process with (right) representative sample images at critical points during fabrication. The optical microscope images at 45° show the status of the electrodes (top) before removal of the sacrificial layer, (center) after removal of the sacrificial layer but before the heating treatment to activate electrode bending, and (bottom) the final state with the vertical PI cantilevers and electrodes.

### Polyimide cantilevers release and bend

To release PI structures, the patterned samples in step F in Fig. 2) were immersed in Technietch Al80, removing the sacrificial layer. Afterwards, the samples were cleaned with deionized water, rinsed with isopropyl alcohol and dried in air, facing down. The samples were heated to 200 °C to soften the PI,^24^ allowing the stress mismatch between SiO_2_ and PI to impose a curvature on the hinge region leading to the vertical structures seen in Fig. 1b and Fig. 2. Further temperature variations do not cause the structures to revert to a planar configuration.

To obtain images of the 3D structures, the samples were placed under a microscope at 45° and several photographs with different focus planes were taken. These were then digitally combined to achieve a single image of the complete sample.

### Electrical characterization

Electrical characterization was performed on a global per-sample scale with Electrical Impedance Spectroscopy (EIS), using an Impedance Analyzer 262k-1000 (SinePhase Instruments), and on a per-electrode level using impedance and noise level mapping, with a MEA-IT60 impedance testing device (Multi Channel Systems MCS GmbH) and miniMEA-2100 system (Multi Channel Systems MCS GmbH), respectively. In all measurements, phosphate buffered saline (PBS) was used as the medium. Before electrical measurements and growth of neuronal cultures, the samples were treated with air plasma in a PIE Scientific Tergeo Plasma Cleaner, with an air flow of 10 sccm and an applied power of 15 W.

EIS is a non-destructive characterization method to measure the frequency-dependent impedance of the electrode array.^25^ EIS for all electrodes was performed by connecting all contact pads on the sample border in parallel using a conductive ink, excluding the reference electrode. The measurement frequency was swept from 1 to 100 kHz to cover the frequencies relevant in electrophysiology.^26^ Impedance mapping was done at a fixed frequency of 1 kHz.

Noise level acquisitions were performed by acquiring voltage fluctuations and using the root mean square (RMS) of the noise level over the duration of the measurement to quantify the performance of each individual electrode. A 200 Hz high-pass filter was used to remove low-frequency fluctuations, including the 50 Hz noise from the electrical grid. Data was acquired at a sampling rate of 20 kHz for 5 min. Measurements acquisition was done using a dedicated software (*Multi Channel Experimenter*). The data was extracted using *Multi Channel Analyzer* and analyzed with a Python script.

### MEAs preparation and neuronal culture

Before neuronal culture growth, planar MEAs were subjected to air plasma treatment and standard coating procedures with poly-D-lysine (20 µg mL^−1^) as described elsewhere.^27^ For 3D MEAs, no air plasma treatment or coating procedures were performed, with the surface hydrophobicity ensuring that the 3D hydrogel maintained a dome shape that covered all electrodes.

Dissociated hippocampal neurons were obtained from Wistar embryo rats (E18). The 3D culture was performed by mixing Geltrex^™^ matrix (A1413301, LDEV-Free, hESC-Qualified, Reduced Growth Factor Basement Membrane Matrix, ThermoFisher Scientific) and a neuronal cell suspension (100 thousand cells per µL) at a 1:3 ratio in cold conditions. A total of 60 µL was pipetted into the central region where the electrodes are located and allowed to settle for 1 min. The device was then incubated upside down at 37 °C for 30 min to promote the polymerization of the hydrogel and the homogeneous distribution of the neurons along 3D spatial arrangements. Finally, the delimiting well was filled with cell culture medium (500 µL per well) and stored in a humidified incubator at 37 °C supplied with 5% CO_2_. Neurons were cultured in Neurobasal Plus medium supplemented with 0.5 mM glutamine, 2% (v/v) B27 Plus, and 1% (v/v) penicillin/streptomycin (P/S), and half-medium changes were performed every 3 days.

### Neuronal activity recordings

The miniMEA-2100 system was used to record the voltage activity of the neuronal culture. Data was acquired at a sampling rate of 20 kHz for 5 min and a temperature of 37 °C was maintained via an external temperature controller. The dedicated software was used to control data acquisition, explore acquired data, and extract the location of neuronal activity spikes. A 200 Hz high-pass filter was used. After the experiments, the cell culture medium was carefully replaced with a 1% solution of the enzymatic detergent Terg-a-zyme^®^ (Z273287, Sigma-Aldrich) and left overnight at room temperature. After repeated washings with distilled water, the samples could be reused without damage to the 3D electrodes.

## RESULTS AND DISCUSSION

### 3D MEAs design

The design of the MEAs was chosen to be compatible with a commercial 60MEA design by Multi Channel Systems MCS GmbH, including the number, size, and arrangement of electrodes, the layout of the pads for external connections and the planar 1.3 × 2.2 mm^2^ reference electrode. The recording electrodes were distributed across 12 PI cantilevers, each 715 µm long and 260 µm wide, within a central circle with a diameter of 6 mm, defined and delimited by the well to confine the neuronal cultures. Each of the PI cantilevers has 5 recording electrodes spaced 100 µm apart along the *z* axis (Fig. 1b, right). The first electrode is located at a height of approximately 150 µm from the substrate, outside the curved region imposed by the SiO_2_ hinge. Images of the final device after the complete fabrication process are presented in Fig. 1b and Fig. 2.

### Stress actuation and controlled vertical angle

PI cantilevers with electrodes were lifted by the stress mismatch between PI and SiO_2_. To ensure precise control of the bending angle of the cantilevers, we tuned the thickness of the thin films and the length of the SiO_2_ layer. The latter acts as a micrometric hinge, allowing us to achieve bending angles close to 90° (Fig. 1b, center), required for the 3D MEAs. This strategy for reaching vertical electrodes is implemented without increasing the complexity of the fabrication process.^16^

PI and SiO_2_ were chosen as the two layers due to their biocompatibility and high resistivity,^16,23^ with both materials having previously been used in similar applications.^9,19^ An analytical model was used taking into account plane strain conditions to assess the effect of the thickness of each material on the curvature of the multilayer (see Supplementary Information and Eq. S1).^15^ The thickness of PI was set to two 3.3 µm layers, as thin as possible while still allowing PI to act as a support and passivation for the electrical leads, and the thickness of SiO_2_ was established at 200 nm resulting in a curvature radius (*R*) of 232 µm.

We then optimized the length of the SiO_2_ micro-hinge to fabricate vertical PI cantilevers. To understand the dependence of the bending angle on the micro-hinge dimensions, multiple PI cantilevers were fabricated with various hinge lengths. Figures 3a-b show two examples of lifted PI cantilevers with 200 and 230 µm hinges, from which the corresponding bending angle was extracted. More details, including additional images with all varied conditions, can be found in Supplementary Information.

**Figure 3:**
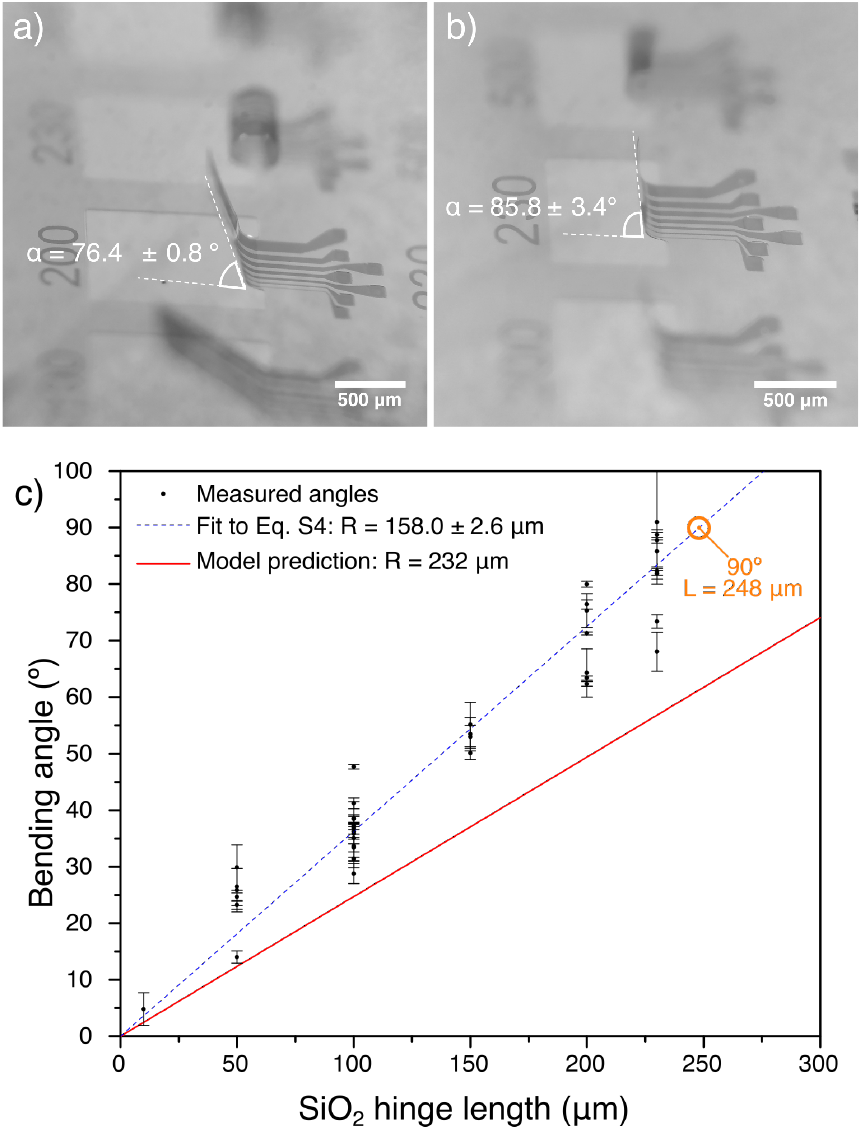
Optical microscope images of a PI cantilever (two 3.3 µm PI layers) and a 200 nm thick SiO_2_ micro-hinge showing: a) an angle of 76.4 ± 0.8°, achieved with a hinge length of 200 µm, and b) an angle of 85.8 ± 3.4°, achieved with a hinge length of 230 µm. c) Bending angle measured between the PI cantilever and the substrate as a function of the SiO_2_ micro-hinge length. The red line shows the relation between angle and hinge length with *R* = 232 um, as predicted by the analytical model. The dashed blue line is a fit to the experimental data points using Eq. S4 and yielding *R* = 158.0 ± 2.6 um. For an angle of 90° we used *L* = 248 µm, highlighted in orange.

Figure S3 shows the experimental data from all the fabricated samples. The calculations from the analytical model do not overlap with our experimental data, and result in an overestimated *R* of 232 µm, which leads to a bending angle smaller than that observed in our samples. This can be explained by the fact that the plane strain approximation does not perfectly describe multilayers with finite lengths. Furthermore, the effects of Ti/Au in the curvature estimation were not considered because of the small thicknesses involved. Fitting the experimental data in Fig. 3c to Eq. S4, we obtain a *R* of 158.0 ± 2.6 µm, which is used to determine the SiO_2_ hinge length *L* required to fabricate a vertical PI cantilever, resulting in 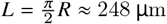.

The wide range of bending angles achieved by stress actuation (Fig. S3) demonstrates the versatility of this technique. Furthermore, the use of a SiO_2_ layer on the entire bottom surface of the cantilever induces the PI to roll into a tubular shape, as shown in some cases of Fig. S2, corresponding to angles exceeding 90°. Such self-rolling structures with embedded electrodes have been employed to encapsulate and record the activity of spherical organoids.^28^ Finally, the electrical resistance before and after stress actuation was evaluated, with no significant changes measured (see Supplementary Information section S3 and Fig. S4).

The residual stresses of each layer play a crucial role in achieving the desired bending angle. These parameters depend on the deposition temperature^29^ or the base pressure of the chamber^16^ and can result in variations among different batches of samples. Nevertheless, we observed that most cantilever structures within a single MEA exhibit similar bending angles, as illustrated in Fig. S5. In fact, the MEAs shown in Fig. 2 are an example in which 100% of the fabricated cantilevers have an angle of 90° relative to the substrate.

Unlike other stress-actuated MEAs that use rigid materials such as SiO_2_/Si_3_N_4_,^19^ in our approach heating is required after the removal of the sacrificial layer. This heating step temporarily softens the PI, allowing the micro-hinge to act, making the cantilever achieve the targeted bending angle.^24^ However, the temperatures used during this step are too high for cells to survive, so performing actuation with neuronal cultures already on the MEAs is not possible. In our samples, there is no mechanism to recover the original planar configuration apart from direct mechanical actuation. However, there is significant interest in having a reversible bending state, as it suits potential applications in organoid envelopment^28^ or cell gripping^30^. Reversible actuation in real time using similar structures could be based on hydrogel swelling^14,31^, or using materials with different thermal expansion coefficients^32^. An alternative could also be manipulation by Joule heating through a patterned region that is thermally and electrically insulated from the culture, similar to results previously reported in literature^16^.

### Electrical characterization of the MEAs

The reported impedance in Ti / Au MEAs is in hundreds of kΩ to a few MΩ range per electrode,^33^ with a limit of 5 MΩ for neuronal recordings. Higher impedance values can lead to voltage noise levels that compromise the ability to accurately capture neuronal signals.^34^ Therefore, to ensure compatibility with the final application, the patterned electrodes (Fig. 4a-b) were characterized in terms of their impedance and voltage noise before and after stress actuation. Figures 4 c-d show the results of the impedance mapping of the Ti / Au conductive layer. For MEAs fabricated following the protocol presented in this work, electrodes with impedances as high as 1.37 MΩ successfully recorded neuronal activity. Using this as a reference for functional electrodes, the average impedance of functioning electrodes at 1 kHz is 0.99 ± 0.22 MΩ for the planar configuration and 0.96 ± 0.07 MΩ for the vertical ones. Figure 4c shows a comparison between the EIS spectrum of a planar MEA (total impedance of 19.9 ± 0.1 kΩ) and a different 3D MEA (23.5 ± 0.5 kΩ). The EIS spectra were acquired with the electrodes connected in parallel. So, assuming that all electrodes have the same impedance, we can estimate per-electrode impedance of 1.17 ± 0.01 MΩ for the planar MEA and 1.39 ± 0.04 MΩ for the 3D MEA. This can be an overestimation since non-functioning electrodes are also included. In general, the results show that the impedance was not significantly altered by the stress actuation process, and the values obtained are within the range reported in literature.^33^

**Figure 4:**
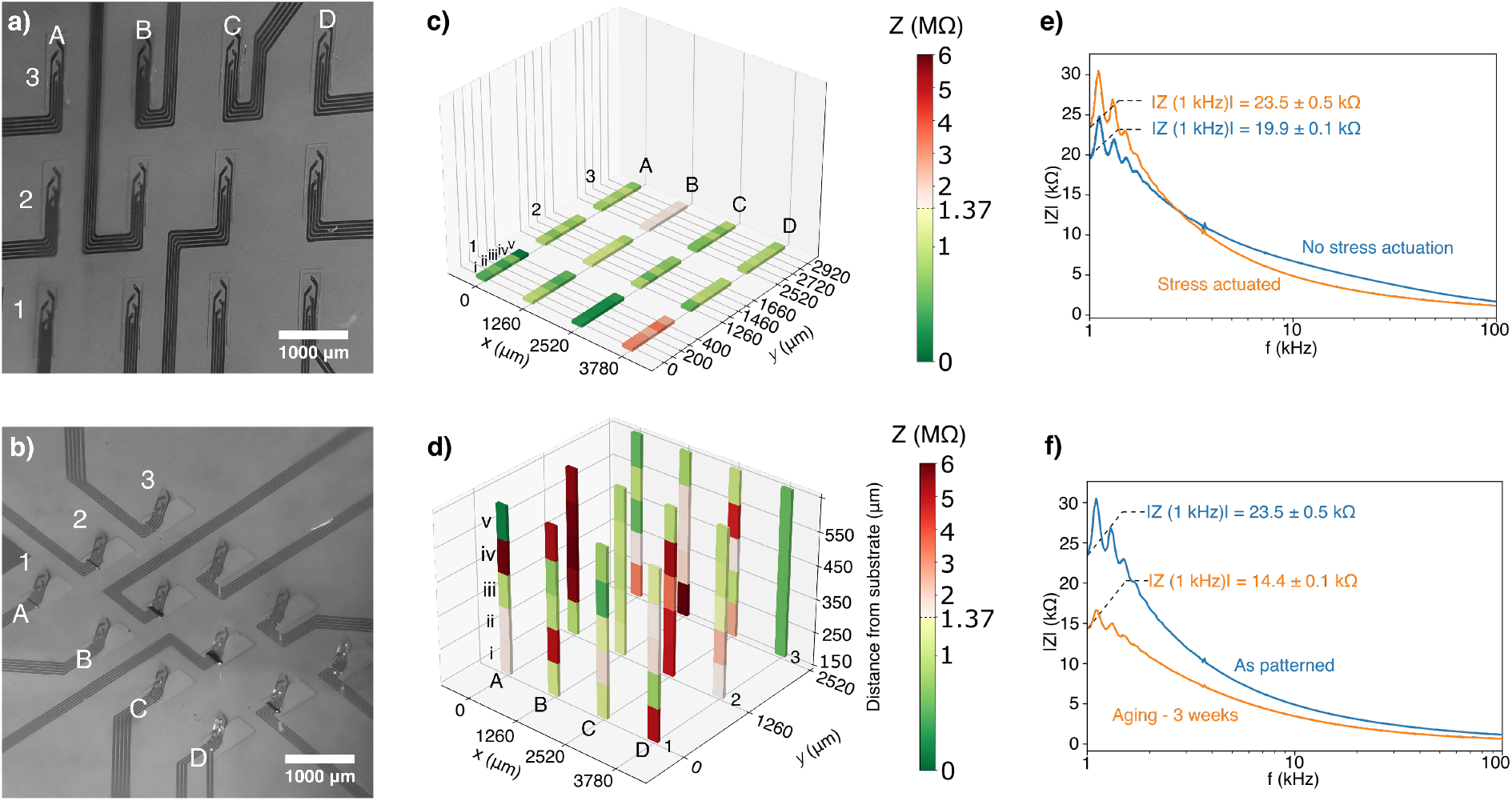
Electrical characterization of the planar and 3D MEAs. Optical image of the measured a) planar MEA and b) 3D MEA. Electrical impedance mapping of c) planar MEA and d) 3D MEA after stress actuation. e) Representative EIS measurements comparing a planar (no stress actuation) and a 3D MEA (stress actuated) highlighting the value of |Z| at 1 kHz for comparison. f) Aging effect on the EIS spectra of a 3D MEA, before and after 3 week immersed in PBS.

One limitation observed in Fig. 4 is the low number of 3D electrodes that are electrically operational. On average, the percentage of functional electrodes is 79% for planar MEAs and 51% for 3D MEAs. This reduced electrical yield can present a challenge for process scale-up. In our experiments, we identified the opening of the via through the PI as the main bottleneck for the decrease in the number of functional electrodes. Fluctuations in the baking times or temperatures of the top PI layer alter the conditions necessary for simultaneous patterning and etching in step E (Fig. 2). This can be mitigated by further optimization of the fabrication process.

The efficacy of PI as an electrical passivation layer was evaluated by measuring the impedance of a patterned MEA with a continuous PI coverage of 3.3 µm. A value of 72.5 ± 5.9 kΩ was measured, more than 3 times higher than in MEAs with vias open through the top PI layer. These results are indicative of the absence of pinholes and minimal current leakage.^35^

Given the extended times needed for neuronal culture growth prior to their characterization, the long-term stability of PI passivation was also assessed. To simulate aging, the samples were left in contact with PBS at 37 °C for 3 weeks.^36^ MEA samples with opened vias showed a slight decrease in impedance at 1 kHz, from |*Z* (1*KHz*) | = 23.5 ± 0.5 kΩ (1.39 ± 0.16 MΩ per electrode) to |*Z* (1*KHz*) | = 14.4 ± 0.1 kΩ (0.85 ± 0.01 MΩ per electrode), as seen in Fig. 4f. The decrease in impedance could originate from the higher hydrophilicity of the leads after 3 weeks immersed in PBS and shows that there was no significant degradation in the conductivity of the Ti/Au leads. For MEAs without opened vias, the same experiment yielded a decrease in impedance from |*Z* (1*KHz*)| = 57.6 ± 2.7 kΩ (3.40 ± 0.16 MΩ per electrode) to |*Z* (1*KHz*) | = 38.3 ± 0.6 kΩ (2.26 ± 0.04 MΩ per electrode), The latter is still higher than the value obtained in electrode arrays with open vias, showing stable passivation. Although other protection strategies could be considered,^37^ any additional material added to the multilayer will affect the stress in the thin films and therefore require different dimensioning of the PI cantilever and SiO_2_ micro-hinge.

The voltage noise level in each electrode was mapped in both planar and 3D MEAs immersed in PBS (see Fig. S7). The electrical noise measured in the fabricated MEAs was independent of frequency in the measured range of 200 Hz to 4 kHz, chosen to include the 1 kHz frequency relevant to electrophysiology.^26^ This is consistent with the dominant thermal noise.^33,34^ Planar MEAs showed average voltage noise levels of 6.93 µV_RMS_, with a standard deviation of 1.36 µV_RMS_. Noise levels in 3D samples skew slightly higher than those of the planar samples, with an average of 7.83 µV_RMS_ and a standard deviation of 1.74 µV_RMS_. In our experiments, we detected neuronal activity with electrodes that had noise levels as high as 9.38 µV_RMS_. Therefore, noise level mapping can be used as a qualification tool to quantify the number of electrodes suitable for recording neuronal activity prior to culture growth.

### Neuronal recordings using 3D MEAs

Neuronal recordings were performed on 3D MEAs to validate the final device application and the fabrication process implemented. The neuronal culture was grown in a hydrogel to ensure that the cells maintained a 3D spatial arrangement. A 5-minute recording of spontaneous activity at 12 days *in vitro* (DIV) is presented in Fig. 5. The MEA used for this purpose is shown in Fig. 1b.

**Figure 5:**
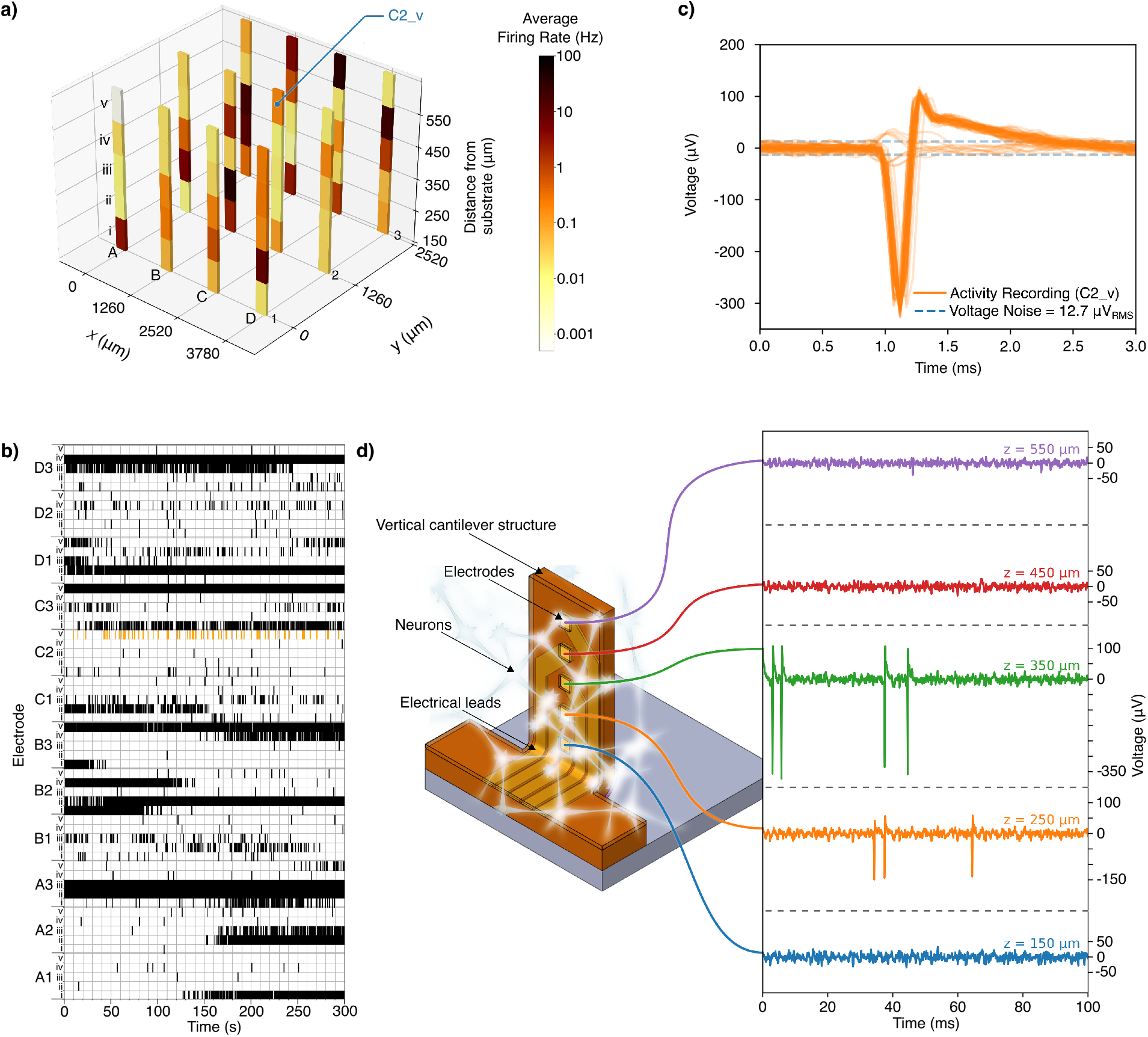
Neuronal activity recording with a 3D MEA at 12 DIV. a) Map of spike activity in a 5 minute recording of spontaneous neuronal activity using a 3D MEA. b) Raster plot of the recording shown in a). Each line represents a single detected spiking event, with clusters of detected spikes showing bursting behavior. Associated with each PI cantilever are the 5 electrodes, each at a specific height. c) Overlay of neuronal activity spikes detected in a 5 minute recording of spontaneous neuronal activity in electrode C2_v in a) and b), at a height of 550 µm above the substrate. d) Simultaneous neuronal activity recordings for electrodes at different heights along a single PI cantilever (A3 in a) and b)). This analysis allows examination of neuronal signal propagation along the *x, y* and *z* axes, demonstrating the viability of these MEAs to study neuronal signals in 3D cultures.

The neuronal spike activity in the sample is shown through the map in Fig. 5a and the raster plot in Fig. 5b. Most electrodes detect some degree of neuronal activity, being significantly more intense in a few select electrodes that present burst behavior. The electrodes with the most activity recorded are distributed throughout the entire culture area, not clustering in a specific region. Electrodes at all heights were able to measure neuronal spikes, validating that the fabricated samples can be used for neuronal activity recording in 3D cultures.

In Fig. 5c, the neuronal activity spikes recorded with a single electrode during a 5-min acquisition of spontaneous activity are overlaid. Peaks with both positive and negative polarity are detected. The polarity of the neuronal activity spike has been shown to correlate with signals being generated at different locations of the cell, or captured at different distances from the signal origin.^38,39^

For reference, neuronal recordings were also performed on planar MEAs (Fig. S8). Given the thickness of the Ti/Au layers, the electrodes are slightly transparent and cells can be seen through them even with bottom illumination, as observed in Fig. S8a. The density of cells was significantly increased in 3D cultures compared to the one used in 2D cultures, to ensure that enough cells were located next to the electrodes in the 3D space to maximize the probability of capturing neuronal activity. Spikes with amplitudes as high as 400 µV were detected with the 3D MEAs, with individual electrodes recording up to 12078 spikes in a 5-minute period, corresponding to an average firing rate of 40.26 Hz. In the same duration, with planar MEAs, spikes of the order of 40 µV were recorded, with a maximum of 1046 spikes per electrode, an average firing rate of 3.49 Hz.

As an example of the capabilities enabled by the MEAs described here, a short recording of electrical activity of the neuronal culture is presented in Fig. 5d for all electrodes in a single vertical structure. Although a complete analysis of cellular dynamics is beyond the scope of this work, the temporal coincidence of spikes in various electrodes can point to a common origin in the cellular culture. This analysis highlights the potential of this MEA platform in conducting studies of signal propagation in neuronal networks along the *x, y* and *z* axes.

## CONCLUSIONS

Stress-actuated 3D MEAs were fabricated, characterized, and used to record neuronal activity, demonstrating the validity of these platforms for electrophysiological studies. The devices produced through our patterning methods have custom layouts, have potential for scaled-up fabrication, and are compatible with commonly used neuronal activity recording setups and protocols. Our proposed 3D MEAs can therefore provide innovative and cost-effective solutions for recording activity in 3D neuronal networks, aiming at driving future developments in neuroscience and electrophysiology.

The performance of the stress-actuated 3D MEAs was comparable to planar MEAs fabricated using similar process steps. The average impedance of the 30 × 30 µm^2^ electrodes in the 3D MEAs was 0.96 ± 0.07 MΩ, within the ranges reported in similar planar structures^33^ and noise levels of 7.83 µV_RMS_ were achieved. These specifications were suitable for recording neuronal activity in 3D, with electrodes at different heights ranging from 150 to 550 µm. Spikes with amplitudes of up to 400 µV and firing rates of up to 40 Hz were detected, in line with other reported solutions that show limitations in fabrication scale-up.^9^

In addition to manipulating the vertical PI cantilevers developed in this work, stress actuation methods can be adapted to fabricate arbitrarily shaped geometries combining different bending angles and directions within a single MEA to obtain customized 3D geometries.^15^ A simple analytical model that approximates the patterned structures to a single bilayer was used as a guide for the design. However, to obtain better estimates, one needs to take into account the Ti / Au layers and allow for the use of compact passivation strategies, such as combining SiO_2_ and Si_3_N_4_^5^,^40^ or using silicone-based gels.^41^

Using PI as material for the stress-actuated cantilevers is advantageous due to its flexibility. Although similar approaches have been reported using rigid materials for organoid penetration,^19^ the use of a flexible polymer ensures Young’s modulus compatibility with neuronal cultures, resulting in enhanced biocompatibility.^20^ Rigid structures are also brittle, making flexible materials more resistant to deformations during handling or neuronal culture growth.^42^ To further enhance the capabilities of flexible 3D MEAs, the neuron-electrode coupling can be improved by using recording electrodes with a protruding shape,^7^ also compatible with the patterning methods employed.

Finally, the versatility of stress actuation allows for the fabrication of customized bendable structures, which could be an advantage when recording electrical activity of cultures with engineered distributions achieved by cell culture patterning^43^ or 3D bioprinting.^44^ Furthermore, 3D MEAs can be extended to other electroactive cellular systems, including cardiac cells^45^ or muscle tissue,^26^, or even be embedded in microfluidic platforms,^22^ opening new directions for future research.

## Notes

### Competing Interest Statement

The authors have declared no competing interest.

