## Supplementary Information for "Stress-actuated Flexible Microelectrode Arrays for Activity Recording in 3D Neuronal Cultures"

### S1. Stress actuation model

We consider a bilayer structure in plane strain conditions (wide strips with bending constraint in one direction).[1] Each of the materials in this layer has thickness $t_i$, Young's Modulus $E_i$, Poisson's ratio $\nu_i$, and is deposited with an initial strain $\varepsilon_i^0$, where $i = 1$ for the bottom material and $i = 2$ for the top material, as represented in **Fig. S1a**. In these conditions,[2] the curvature $K = 1/R$ (where $R$ is the radius of curvature) is given by:

$$K = -\frac{6E_1'E_2't_1t_2(t_1+t_2)(\eta_1\varepsilon_1^0 - \eta_2\varepsilon_2^0)}{(E_1')^2t_1^4 + (E_2')^2t_2^4 + 2E_1'E_2't_1t_2(2t_1^2 + 2t_2^2 + 3t_1t_2)}, \qquad (S1)$$

where the variables $E_i'$ and $\eta_i$ are defined as:

$$E_i' = \frac{E_i}{1-\nu_i^2}, \qquad (S2)$$

$$\eta_i = 1 + \nu_i. \qquad (S3)$$

In Eq. S1, the sign represents the direction of curvature, with $K$ being positive when the multilayer bends towards the top layer, away from the substrate.

As main materials for our MEAs design we consider:

a) Spin-coated PI layer as the top material ($E_2 = 245$ GPa, $\nu_2 = 0.34$,[2] $\sigma_2 = E_2\,\varepsilon_2^0 = 3.6$ MPa)[3] and
b) $SiO_2$ layer deposited by magnetron sputtering as the bottom material ($E_1 = 77$ GPa, $\nu_1 = 0.202$, $\sigma_1 = E_1\,\varepsilon_1^0 = -300$ MPa),[2]

The radius of curvature can then be plotted as a function of the thickness of the $SiO_2$, for multiple thicknesses of PI, as shown in **Fig. S1b**. The chosen PI thickness imposes a minimum radius of curvature; as such, the thickness of the PI layer was chosen to be as thin as possible to ensure the desired flexibility for bending (6.6 μm, obtained with two 3.3 μm coatings), allowing control over the angles that can be achieved using stress actuation. The $SiO_2$ thickness was set at 200 nm, achieving a radius of curvature of $R = 232$ μm, which translates to sharp angles, without excessive deposition time.

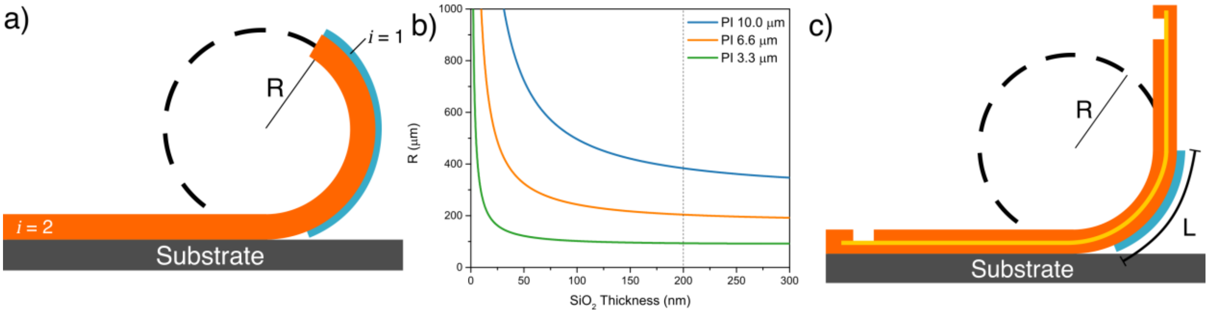

*Figure S1 – Stress-induced curvature model to obtain self-assembled vertical arrays of electrodes. a) Schematic representation of the modelled bilayer, with a stress-inducing layer (blue layer) forcing a curvature on the PI (orange layer). b) Plot of the radius of curvature of a PI/$SiO_2$ bilayer for various PI thicknesses, as a function of the $SiO_2$ thickness. c) Vertical cantilevers obtained by patterning the $SiO_2$ layer into a hinge with length L.*

### S2. Controlling the bending angle

Patterning the $SiO_2$ layer into a limited hinge region, as shown in **Fig. S1c**, forces the PI cantilevers to bend upwards at a specific angle set by the hinge length $L$. The angle $\alpha$ (in radians) between the cantilever and the substrate - after stress actuation - can be calculated from the geometric definition for the arc length in a circle with radius $R$:

$$\alpha = \frac{L}{R}. \qquad (S4)$$

By changing $L$, we are able to control the bending angle, as demonstrated in **Fig. S2**. There, we show the possible bending angles of the PI cantilevers as a function of both the cantilever length and the hinge length. In **Fig S2**, the length of the PI cantilever is changed between 100 μm and 2000 μm (vertical axis), while the length of the $SiO_2$ hinge ($L$ in Eq. S4) is varied between 0 to 2000 um (horizontal axis). For the latter case, several repetitions of patterned structures were also included for reproducibility assessment. Making the $SiO_2$ layer cover the entire PI cantilever structure leads to a self-rolling effect, which results in the tubular structures seen in the bottom right of **Fig. S2**.

Figure 3 of the manuscript shows the dependence of the bending angle on the hinge length $L$ focused on the targeted vertical geometry at 90º. **Figure S3** complements Fig. 3 by plotting all fabricated lengths, including those that lead to bending angles above 90º and even rolled structures with angles above 360º. The plots in Fig. 3 and **Fig. S3** include bending angles for cantilevers with different lengths. Here, the influence of the PI cantilever length on the bending angle is neglected, as no PI curvature was observed in the sections outside the hinge structures.

From **Fig. S3**, the hinge length required for 90º structures is 248 μm, which is the value used for the final MEAs design. The PI cantilever length (715 μm) was chosen to accommodate the micro-hinge and 5 electrodes separated by 100 μm. The width was set at 260 μm to fit 30 μm-wide leads for access to all 5 electrodes with good electrical conductivity, in addition to tolerances for the PI patterning process.

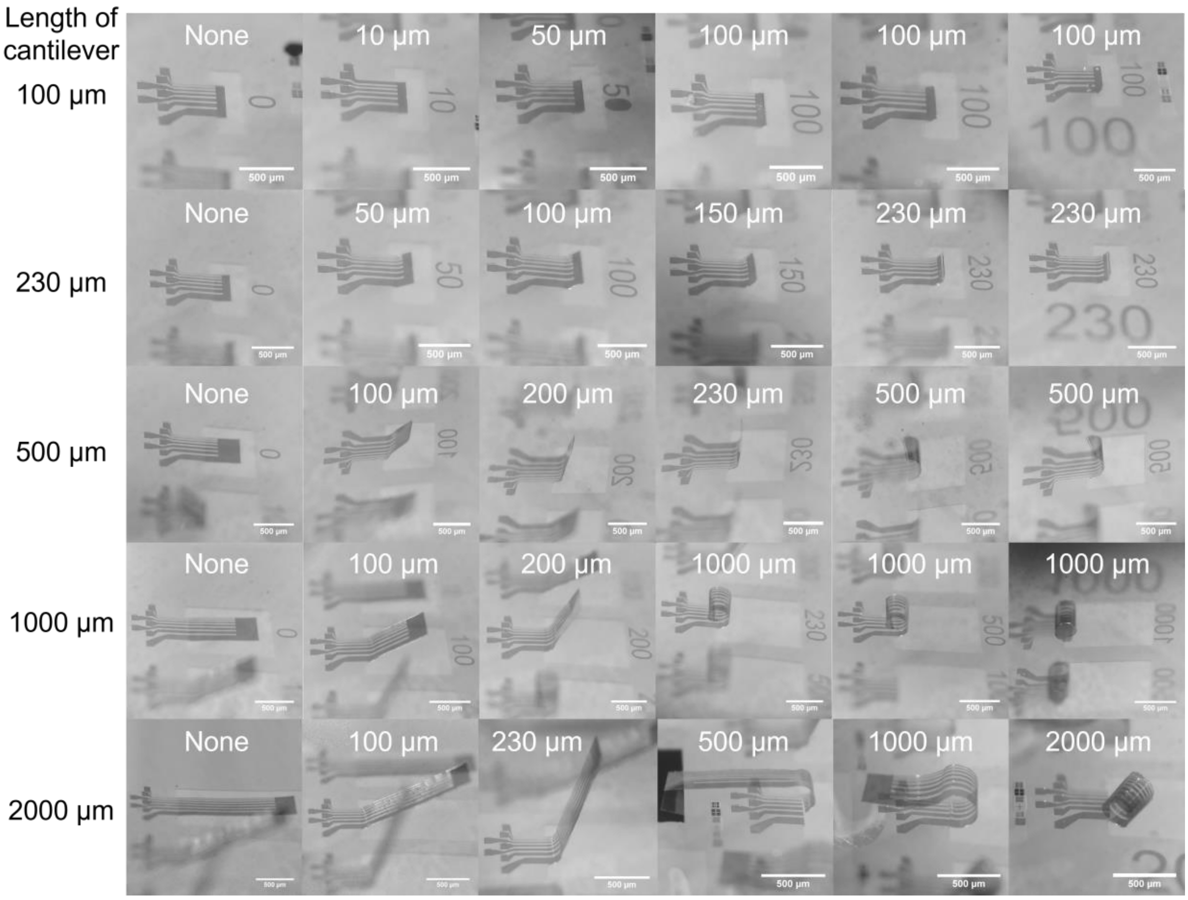

*Figure S2 – PI cantilevers with different lengths and with different $SiO_2$ micro-hinge lengths, showing the possibility of obtaining multiple bending angles and, as such, different electrode heights in a single sample. The text in each image indicates $L$ - the length of the hinge. On the left side is the corresponding PI cantilever length.*

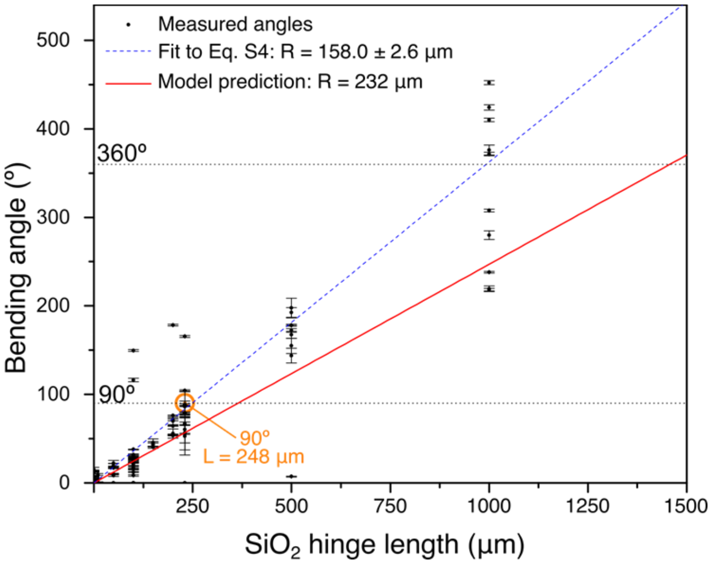

*Figure S3 – Measured bending angle of two 3.3 µm PI layers as a function of the length of a 200 nm $SiO_2$ thick micro-hinge. Angles above 360°, obtained in hinges larger 1000 µm, represent samples with a rolled geometry. The red line shows the prediction of the plane strain model (Eq. S1) using input parameters for PI and $SiO_2$ (see section S1) yielding $R = 232$ µm. The blue dashed line is a fit to the experimental data using Eq. S4, yielding $R = 158.0 \pm 2.6$ µm. The hinge length required for a 90º bend is highlighted in orange. The radii of curvature and angles were measured from microscope images seen from the side.*

### S3. Impact of bending on the Ti/Au leads electrical properties

Test samples with 2000 µm-long, 500 µm-wide PI cantilevers each containing 50 µm-wide 5 nm Ti / 20 nm Au leads were characterized. The V(I) of the leads was performed in a planar configuration (**Fig. S4a**) corresponding to the blue curve in **Fig. S4c**, and then in a rolled geometry (**Fig. S4b**) corresponding to the orange curve. The rolled geometry features stress actuation across the entire length of the leads, therefore maximizing any potential performance degradation.
The resistances of the leads are basically unaltered, with an average difference of only ΔR = 1.1 ± 33.1 Ω (a change of around 1.4% ± 15.8%) in 3 measured samples. This is expected, as the conductive layer is positioned close to the neutral plane.[4,5]

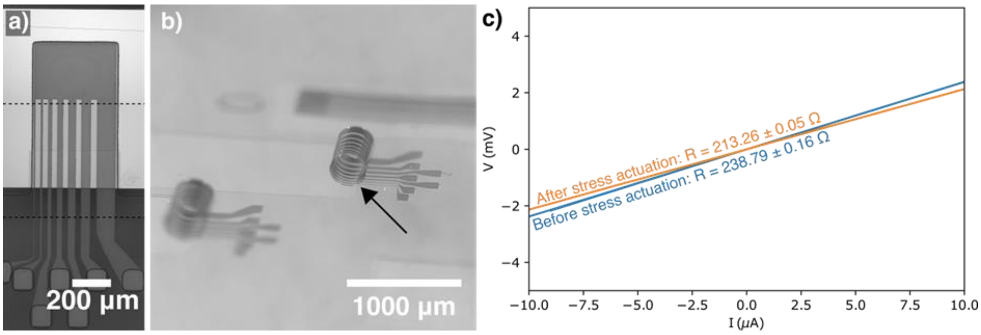

*Figure S4 – Electrical resistance characterization before and after stress actuation. a) Microscope image of a planar PI cantilever in the fabricated test samples This image was composed of multiple photographs of the same structure joined at the dashed lines. b) Microscope image of a different structure in a rolled geometry. c) Representative I-V measurement before and after stress actuation for the structure highlighted in b). The average resistance increase due to rolling for similar samples is of* $\Delta R = 1.1 \pm 33.1\ \Omega$.

### S4. Reliability of stress actuation

The fabrication process described in the manuscript results in uniform bending angles across multiple structures within a single sample. This is highlighted by **Fig. S5**, where images of multiple samples are shown, showing overall uniform near-vertical structures within the same sample.
We attribute the differences in the bending angle among all distinct samples to variations[2] in the residual stress imposed to the critical layers as discussed in the manuscript (Section *Stress actuation and controlled vertical angle*).

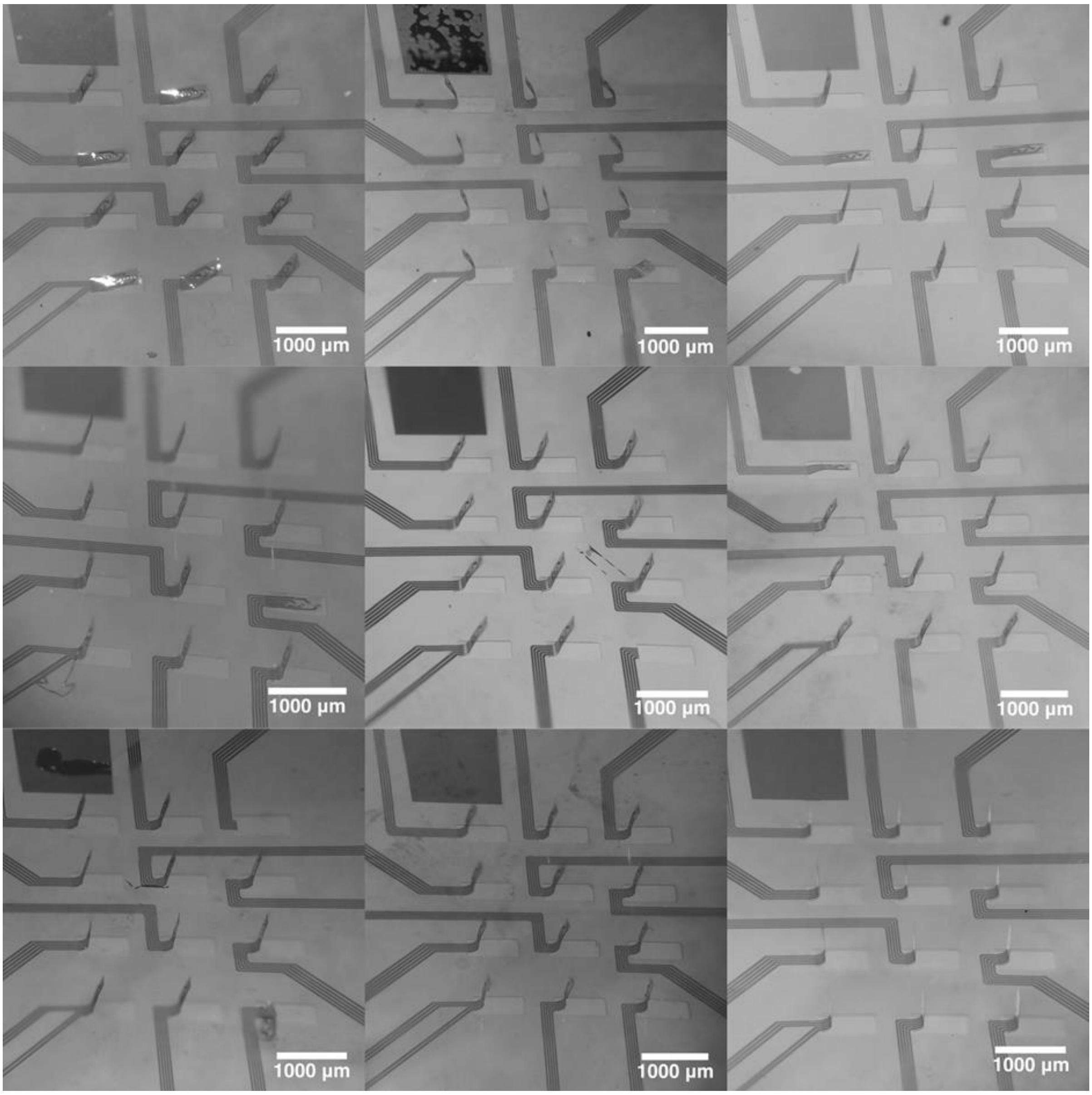

*Figure S5 – Examples of 9 patterned 3D MEAs samples with stress actuation.*

### S5. Additional electrical characterization of patterned samples

PI is not only used as flexible layer to bend but also as a protection layer for the electrical Ti/Au leads.[6] Therefore, we assessed the efficiency of the 3.3 μm PI layer in preventing shunting between the electrical leads and the saline medium.

**Figure S6** shows the spectra of two planar MEA samples, one covered with a continuous PI layer (i.e. electrodes completely encapsulated by the PI layers), and another with vias open through the top PI layer enabling electrical contact to the recording electrodes.

The sample with a continuous PI layer shows an impedance of 72.5 ± 4.9 kΩ (corresponding to 4.28 ± 0.29 MΩ per electrode), much higher than the values obtained when the electrodes are in contact with the solution (18.7 ± 0.1 kΩ, or 1.11 ± 0.01 MΩ per electrode), thus validating the choice of PI as a passivation layer.

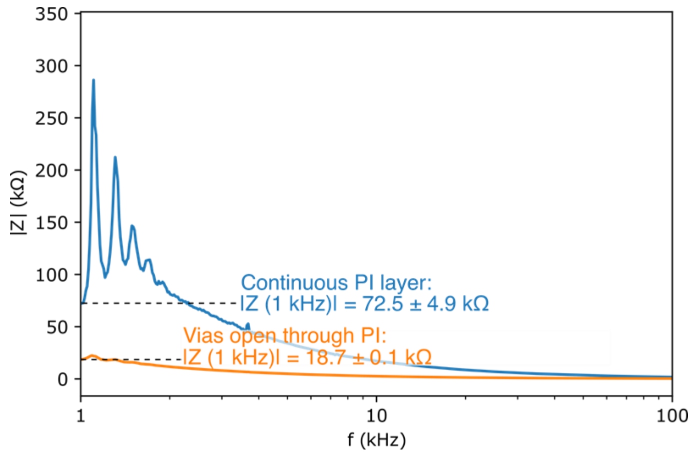

*Figure S6 – PI layer evaluation as suitable passivation for the electrical leads using EIS. The average impedance of samples with continuous PI layer at 1 kHz is of* $72.5 \pm 4.9$ *kΩ, while for samples with vias opened through the PI layer the average impedance is of* $18.7 \pm 0.1$ *kΩ*

Noise level analysis was performed in planar and 3D MEAs. Maps of the voltage noise levels of each electrode in these samples are presented in **Fig. S7**. Average noise levels of 6.93 $\mu V_{RMS}$ and of 7.83 $\mu V_{RMS}$ were obtained for planar and 3D MEAs, respectively. Despite the noise levels in 3D MEAs being slightly higher, both types of electrodes presented low enough values for neuronal activity detection. Neuronal activity spikes were detected with electrodes showing voltage noise as high as 9.38 $\mu V_{RMS}$.

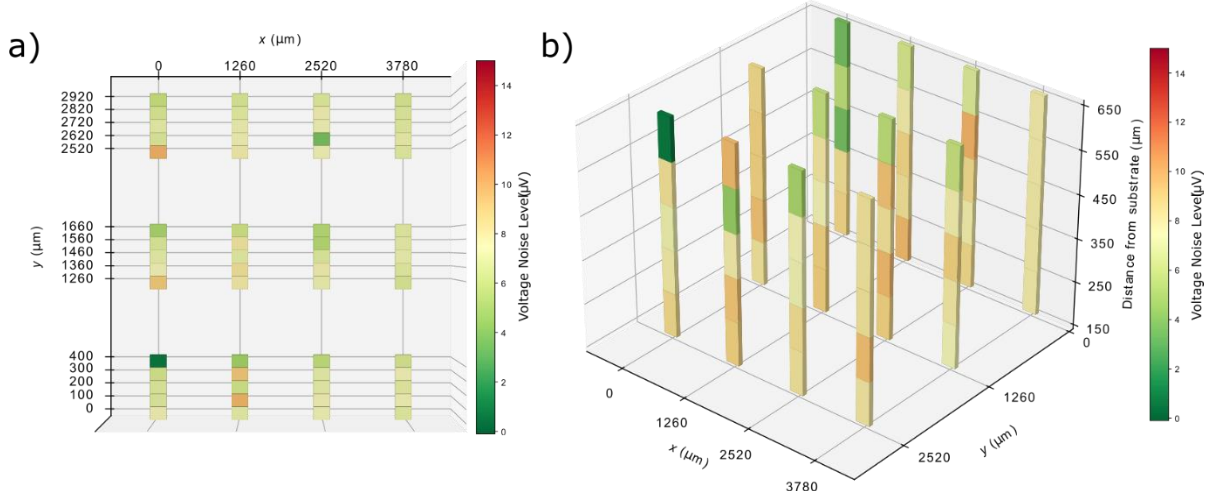

*Figure S7: Voltage noise level map for representative samples with electrodes in (a) planar and (b)3D MEAs.*

### S6. Neuronal activity recording with planar MEAs

For reference, neuronal activity recordings were also performed with planar MEAs, with results presented in **Fig. S8**. The intensity and amplitude of the spikes in **Figs. S8b - d** is lower in this recording than in the recording shown in Fig. 5 in the manuscript. This is expected since the 3D culture was prepared with a higher density of neurons, to ensure that signals would be detected despite the increased volume. The results for 3D MEAs are comparable to those achieved with planar MEAs, demonstrating the viability of the platforms developed in this work.

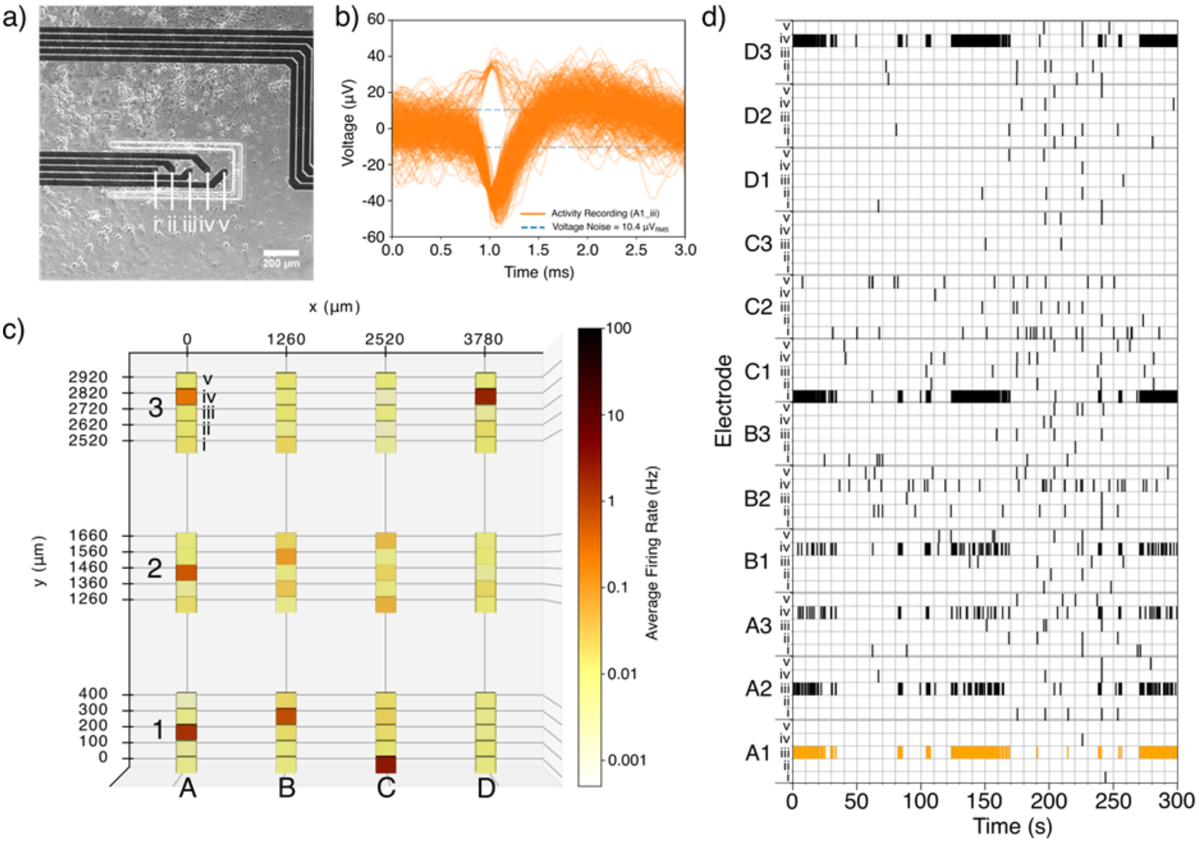

*Figure S8 – Spontaneous neuronal activity recording with a planar MEA at 20 DIV. a) Optical microscope image of neuronal culture on the sample. The image was acquired from a top view, with bottom illumination. The neurons are visible through the electrodes due to the low Ti / Au thickness used. Within a single PI cantilever, electrodes are identified by a number as shown in this image. b) Overlay of neuronal activity spikes detected during a 5-minute recording in the electrode A1_iii highlighted in orange in d). c) Map of spike activity in the array shown in Fig. 4a of the manuscript. d) Raster plot of a 5-minute recording, using a planar MEA. Each vertical line represents a single detected spiking event. Bursting behavior is observed, shown here as clusters of detected spikes.*

### S7. Data treatment and visualization

Impedance mapping data was acquired using the MEAIT software by Multi Channel Systems GmbH, exported into .txt files and visualized with a custom Python script using the Matplotlib graphing library.

EIS data was acquired using the Sinephase Impedance Analyser software. Each sample was measured four times and exported to four text files. A Python script then calculated the average for each point in the four measurements and plotted the average line.

Noise level measurements and neuronal activity recordings were both performed using the Multi Channel Experimenter software, with the Multi Channel Analyzer software (both Multi Channel Systems GmbH) then being used to export the data into text files. For the noise level acquisitions, a Python script then calculated the voltage root mean square for each electrode. For the neuronal activity recordings, spike locations and frequency were also exported to text files using the dedicated software. Matplotlib was then used to display all the acquired noise level and activity data according to the physical location of our electrodes, as well as plot overlaid neuronal activity spikes.
